# Coverage profile correction of shallow-depth circulating cell-free DNA sequencing via multi-distance learning

**DOI:** 10.1101/737148

**Authors:** Nicholas B. Larson, Melissa C. Larson, Jie Na, Carlos Sosa, Chen Wang, Jean-Pierre Kocher, Ross Rowsey

## Abstract

Shallow-depth whole-genome sequencing (WGS) of circulating cell-free DNA (ccfDNA) is a popular approach for non-invasive genomic screening assays, including liquid biopsy for early detection of invasive tumors as well as non-invasive prenatal screening (NIPS) for common fetal trisomies. In contrast to nuclear DNA WGS, ccfDNA WGS exhibits extensive inter- and intra-sample coverage variability that is not fully explained by typical sources of variation in WGS, such as GC content. This variability may inflate false positive and false negative screening rates of copy-number alterations and aneuploidy, particularly if these features are present at a relatively low proportion of total sequenced content. Herein, we propose an empirically-driven coverage correction strategy that leverages prior annotation information in a multi-distance learning context to improve within-sample coverage profile correction. Specifically, we train a weighted k-nearest neighbors-style method on non-pregnant female donor ccfDNA WGS samples, and apply it to NIPS samples to evaluate coverage profile variability reduction. We additionally characterize improvement in the discrimination of positive fetal trisomy cases relative to normal controls, and compare our results against a more traditional regression-based approach to profile coverage correction based on GC content and mappability. Under cross-validation, performance measures indicated benefit to combining the two feature sets relative to either in isolation. We also observed substantial improvement in coverage profile variability reduction in leave-out clinical NIPS samples, with variability reduced by 26.5-53.5% relative to the standard regression-based method as quantified by median absolute deviation. Finally, we observed improvement discrimination for screening positive trisomy cases reducing ccfDNA WGS coverage variability while additionally improving NIPS trisomy screening assay performance. Overall, our results indicate that machine learning approaches can substantially improve ccfDNA WGS coverage profile correction and downstream analyses.

## 1. Introduction

Circulating cell-free DNA (ccfDNA) is comprised of relatively short fragments of genomic material that naturally occur in bodily fluids and originate primarily from normal cell apoptosis^1^. A number of biomedical applications have been identified for shallow-depth whole-genome sequencing (WGS) of plasma ccfDNA, including liquid biopsy for early identification of invasive tumors^2^ as well as non-invasive prenatal screening (NIPS) for fetal genetic and genomic abnormalities^3,4^. For downstream inference, these data are typically summarized by generating binned coverage profiles of the sequencing output, whereby the genome is uniformly partitioned into moderately sized contiguous regions (e.g., 10-50 kilobases (kb)) and count-based coverage is calculated by the enumerating the overlapping sequencing reads. Evidence of copy-number variants (CNVs) and aneuploidy may be detected from these profiles using standard CNV detection methods for coverage data^5^.

Coverage profile patterns for plasma ccfDNA WGS exhibit a large degree of non-uniformity relative to standard nuclear DNA WGS. This is in part attributable to the fact that there is a biased preservation of DNA originating from regions that are protected from degradation by nucleases in the blood stream, including nucleosome- and protein-bound DNA^6^. These patterns can in turn be exploited via deconvolution to identify evidence of tissue-of-origin admixture, and have been leveraged using machine learning methods to build fetal fraction predictors for NIPS^7^ and improve detection of micro-duplications and deletions in NIPS analyses^8^. However, ccfDNA exhibits a large deal of inter-sample heterogeneity that cannot be fully attributed to these patterns, and GC-content correction only explains a moderate proportion of this variability. In the context of NIPS and fetal trisomy detection, this may necessitate higher fetal-fraction quality control thresholds to achieve desired assay sensitivity and specificity, delaying recommended gestational age for the assay and leading to repeat maternal blood draws when estimated fetal fraction in too low.

In this paper, we explore alternative machine learning strategies for improving within-sample coverage profile correction relative to standard regression-based methods commonly implemented for GC-content correction. Using a large set of plasma ccfDNA WGS coverage profiles from NIPS analyses, we propose and train a k-nearest neighbors (kNN) type of approach to leverage empirical bin-to-bin similarities and further integrate prior knowledge captured in genomic annotation sources via a multi-distance learning framework. We compare coverage profile variability reduction in real NIPS maternal plasma ccfDNA WGS data, and additionally characterize potential improvement in discrimination of trisomy cases and negative controls for common fetal trisomies of 13,18, and 21. Finally, we discuss further research directions in the area of ccfDNA WGS coverage data analysis.

## 2. Methods

### 2.1. Data description

#### 2.1.1. Samples

De-identified samples from research and clinical NIPS results conducted by the Genomics Laboratory at Mayo Clinic were considered eligible for this study. Of these, we identified a total of 476 single fetus normal karyotype pregnancy samples, 145 positive trisomy samples (10 trisomy 13, 41 trisomy 18, 104 trisomy 21), and 790 non-pregnant female donor samples for our analyses. Plasma was obtained from blood and stored in a Streck Cell-Free DNA Blood Collection Tube (Streck, Omaha NE). Use of these data for research purposes was approved by the Institutional Review Board.

#### 2.1.2. Shallow-depth whole-genome sequencing

DNA was extracted from plasma using the Qiagen Circulating Nucleic Acid Kit (Qiagen, Venlo Netherlands) and library preparation was conducted using the Illumina TruSeq® Nano DNA Sample Preparation Kit (Illumina, San Diego CA). Sequencing was performed on the Illumina HiSeq 2500 in Rapid Run mode to generate 50-cycle single-end reads, which were aligned to the hg19 human genome reference using Novoalign (Novocraft, Selangor Malaysia). Chromosomal coverage summaries for 50 kilobase (kb) contiguous genomic windows were generated from the resulting BAM files using the WANDY bioinformatics pipeline (http://bioinformaticstools.mayo.edu/research/wandy/), an in-house developed workflow for bin filtering, GC correction, and normalization of low-depth whole-genome sequencing output to identify copy-number variants and aneuploidy.

#### 2.1.3. Data preprocessing

A total of 57,633 50 kb autosomal genomic bins were initially pre-filtered using an in-house defined set of bins that were previously classified as being unreliable (e.g., poor mappability, repeat regions), resulting in *B* = 49,867 bins (87%) under consideration. Raw coverage values for remaining bins were defined as the number of ccfDNA sequencing reads whose start overlaps each bin. We then normalized coverage values within sample by dividing by the mean coverage value across bins. We designate these intermediate coverage values as the *B* × *N* matrix ***X*** for some *N* set of observed coverage profiles.

### 2.2. Neighborhood-based coverage correction

GC-content-based coverage correction methods can be generally conceptualized as grouping bins by a shared annotation characteristic, such that bin strata defined by the same GC value serve as the basis for removing the bias induced by the sample-specific GC-coverage relationship. However, latent features other than GC content may similarly manifest in empirical patterns of bin coverage through evidence of clustering in retrospective data, and the GC-coverage functional relationship may not be consistent across all bins for a given sample.

We alternatively propose the use of empirical bin-to-bin similarity from retrospective data as a way to improve coverage profile de-noising via a k-nearest-neighbors (kNN) type of approach, which requires the calculation of a *B* × *B* dissimilarity matrix, ***D***. However, due to the high dimensionality of *B*, we may additionally wish to leverage *a priori* knowledge about fixed genomic annotation features (e.g., GC content) which contribute to a large proportion of bin coverage variability. Combining annotation dissimilarity with empirical dissimilarity measures in some supervised fashion is desirable, as the former could impart some form of regularization on empirical coverage dissimilarity measures and improve overall coverage correction performance, particularly if our training data suffers from small *N* dimensionality relative to *B*. This amounts to a multi-distance learning problem^9^, as we are seeking to optimally combine dissimilarities from multiple feature sets as a final input feature representation for model training.

#### 2.2.1. Weighted distance averaging

Consider a distance function that is additive, such as the squared Euclidean distance (SED), whereby the dissimilarity measure between bins *i* and *j* with corresponding feature vectors ***x***_***i***_ and ***x***_***j***_ is defined as

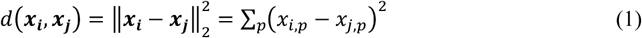

If we have two feature sets ***X*** and ***Y*** from which respective distance matrices ***D***_***X***_ and ***D***_***Y***_ may be calculated using *d*(·,·), these can be efficiently combined under such a distance function by appropriately concatenating and weighting the feature vectors^10^. Define *D*_*α*_(*i, j*) to be element (*i,j*) in a *B* × *B* distance matrix defined as ***D***_*α*_ = (1 − *α*)***D***_*X*_ + *α****D***_***Y***_, where *α* is a mixing parameter that defines the weighted contribution of each component distance matrix. It is simple to show that *D*_*α*_(*i, j*) is equivalent to *d*(*Z*_*i*_, *Z*_*j*_), where 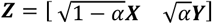 is the weighted column concatenation of matrices ***X*** and ***Y***, since

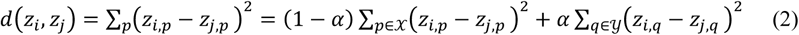

where 𝒳 and 𝒴 denote the respective feature sets unique to ***X*** and ***Y***. This relationship is computationally advantageous, as being able to concatenate the feature spaces in this manner facilitates the use of rapid distance calculation algorithms that identify the leading *k* neighbors and their corresponding distances within a single input feature set, rather than computing the complete distance matrices ***D***_***X***_ and ***D***_***Y***_ prior to weighted combination, which is computationally and memory intensive.

To implement our approach, we applied the kd-tree searching functions for nearest neighbor indices and distances as implemented by the FNN R package, which support fast Euclidean distance calculations for the *k* nearest neighbors via the approximate near neighbors C++ library^11^. Since the relationship between SED and Euclidean distance is monotone, this is equivalent for identifying nearest neighbors under SED (although the output distances can be squared to maintain SED distance values). To also ensure distances are comparable across the feature sets prior to combining, we adopted the double-scaled Euclidean approach for distance normalization, whereby column *p* in ***Z*** is further divided by 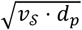, where *v*_*s*_ is the column dimensionality of the respective feature set *S* ∈ {𝒳, 𝒴} to which *p* originally belongs, and *d*_*p*_ is the maximum potential distance for feature *p* as defined by (max_*i*_ (*z*_*i,p*_) −min_*i*_(*z*_*i,p*_))^2^

#### 2.2.2. Genomic annotation

In addition to empirical bin dissimilarity measures derived from coverage profiles, ***X***, we designate the prior annotation information ***Y*** to be comprised of two specific genomic bin annotation sources: (1) GC content and (2) mappability scores, such that ***Y*** is *B* × 2. Both of these have strong *a priori* relationships with coverage profile variability, as depicted in Figure 1.

**Fig. 1.**
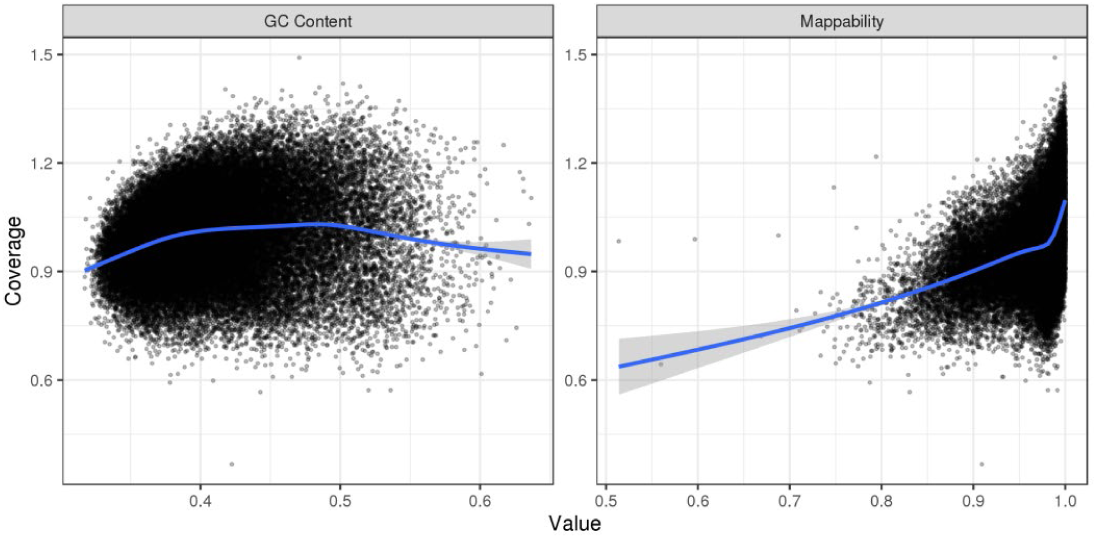
Example coverage relationships with GC content and mappability for a given sample.

### 2.3. Standard annotation-based correction

GC content can heavily influence coverage profiles derived from WGS^12^, and a standard pre-processing step for WGS coverage profile data prior to CNV analysis is within-sample GC-content correction. For a given sample, genomic bins are typically stratified by a shared GC content value and the mean or median stratum-specific coverage value (often with additional lowess smoothing) is used for further normalization. For our coverage correction approach in this manner, we fit generalized additive models (GAMs) using the *gam* R package, where smooth functions are fit using GC content and mappability values as predictors for observed bin coverage for a given sample. Corrected coverage values were then derived by dividing the observed coverage by predicted coverage values produced by the fitted model.

### 2.4. Proposed coverage correction approach

Our basic strategy is to use information in ***D***_***α***_ to define a neighborhood of bins for each candidate bin to serve as the basis for within-sample coverage correction using a k-nearest neighbors (kNN) type approach. Since bins within the physical neighborhood of a given bin (i.e., in *cis*) may also have a higher likelihood of having highly correlated coverage, including these bins in the correction procedure may inadvertently over-correct true copy-number alterations that overlap those bins. This is particularly true in the instances of aneuploidy, where abnormal numbers of chromosomes could be less detectable if neighborhoods were largely comprised of *cis* bins. Thus, we restrict the potential neighboring bins to be those in *trans* with the candidate bin *b*, such that they occur on different chromosomes than *b*. To facilitate this, we split the data by individual chromosomes and their complement, such that bins in the chromosome of interest are queried against the complement for nearest neighbors in the feature space. This also allows for some degree of parallelization and improves overall training computational efficiency.

For coverage correction, we considered both an unweighted and dissimilarity-weighted kNN (wkNN) strategy for within-sample coverage correction, such that raw coverage values of the *k* nearest *trans* bins are used to generate a mean prediction for said bin. That is, for a *B* × 1 vectory ***x*** of raw coverage profile data for a new sample, we define the predicted coverage for bin *i* as 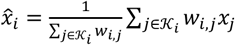 where 𝒦_*i*_ is the size *k* neighboring set for bin *i* and *w*_*i,j*_ is defined as 1/*D*_*α*_(*i,j*) for the distance-weighted approach and *w*_*i,j*_ = 1∀*i,j* in the unweighted version. To perform coverage correction, we again define 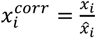, such that the observed value is divided by the predicted value. More generally, the model itself can be represented simply by a *B* × *B* sparse weight matrix ***W*** where *W*_*i,j*_ = *w*_*i,j*_ for 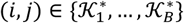 where 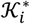 are of tuples corresponding to the query bin *i* and its neighboring bins *𝒦*_*i*_, and 0 otherwise, and 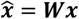

### 2.5. Performance evaluation

To tune the model parameters (*α, k*), we adopted a simple grid search in combination with 5-fold cross-validation in our training set of N = 790 non-pregnant female donor samples. We elected to use only these non-pregnant samples for model training and validation because contaminating fetal ccfDNA in maternal plasma has a different coverage profile due to underlying epigenomic differences^7^. Training a model using data from pregnant female samples could lead to neighborhoods of bins that highly correlate with fetal fraction of ccfDNA and inadvertently over-correct true fetal genomic signal in the coverage data.

Under the cross-validation framework, we considered the samples within ***X*** to be the mode of cross-validation, such that the columns of ***X*** were split into folds for purposes of contributing to ***D***_***α***_ or to performance evaluation. We set the potential tuning parameter values to be *k* ∈ (5, …, 300) at increments of 5 and *α* ∈ (0,0.001,0.01,0.1,0.25,0.5,0.75,1), such that *α* = 0,1 respectively denote models based entirely on ***X*** or ***Y***, respectively. We defined our loss function for tuning as the mean absolute error (MAE) of the individual out-of-fold sample coverage profiles across samples, such that 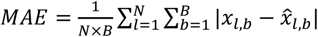 A final model parameterization was then selected based on best cross-validated performance and fit on the complete training data.

For model testing to characterize coverage profile correction performance, we considered N = 476 available clinical NIPS samples ostensibly free of trisomies per previously reported screening results. We considered the within-sample median absolute deviation about the median (MAD), a commonly used metric to characterize sample coverage variability in CNV analysis of WGS data, as a sample-level performance metric for coverage profile correction. As a baseline comparator method for coverage correction, we also performed standard within-sample annotation-based coverage correction described in Section 2.3.

#### 2.5.1. Fetal trisomy detection

In addition to variability reduction, we want to ensure our approach preserves true positive signals at a comparable degree relative to standard coverage profile correction methods. In NIPS, we typically derive chromosome-wise proportions of coverage profiles to determine the presence or absence of common fetal trisomies (i.e., 13, 18, and 21). That is, for a given sample bin coverage vector ***x*** and chromosome *c*, we define proportion 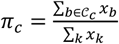 where 𝒞_*c*_ is the set of bins that correspond to chromosome *c* and 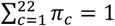. We considered the simple Z-score approach for trisomy detection^13^, such that a large fixed set of reference normal NIPS samples free of trisomies is used to characterize population distributional properties about *π*_*c*_ (i.e., mean *μ*_*c*_ and standard deviation *SD*_*c*_) for trisomy-prone chromosomes. Then, the screening test statistic is defined as

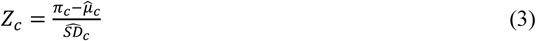

For our purposes, we randomly selected 300 negative NIPS samples to serve as the reference set. Trisomy screening Z-scores were then calculated for the remaining 176 negative samples along with 145 positive trisomy samples (10 Trisomy 13, 41 Trisomy 18, 104 Trisomy 21).

Screening performance for our available NIPS samples already achieves near perfect discrimination using standard coverage profile correction methods due to imposed quality control standards (e.g., minimum sufficient fetal fraction). To assess significant signal improvement (i.e., larger Z-scores for positive cases), we alternatively performed a Wilcoxon signed rank test of the paired Z-scores under each coverage correction approach by trisomy. Evidence of increased Z-scores for positive cases (but not in controls) would indicate that existing thresholds could be relaxed, reducing the number of quality control failures and improving overall assay sensitivity/specificity.

### 2.6. Code availability

All analyses were performed using R version 3.5.2 (R Core Team, Vienna, Austria). Relevant R code is publicly available at https://github.com/nblarson/ccfdna_coverage.git.

## 3. Results

### 3.1. kNN training

The MAE performance measures for the non-pregnant donor data are presented in Figure 2 for the unweighted kNN approach across the considered tuning parameter values. Results were highly comparable to the weighted approach, with a median difference in MAE of 1.0E-05 in favor of the weighted method (range: 0 – 1.8E-04). The optimal tuning parameter settings were also the same for both types of approaches (*α* = 0.01 and *k* = 150), which also led to nearly identical cross-validation performance results (MAE = 0.04134 for both methods). In contrast, using the same *k* = 150, performance for *α* = 0 and *α* = 1 were respectively 0.04137 and 0.06816. Overall, these results indicate that the bins were very densely arranged in the feature space, as the selected *k* is quite large and the weighted kNN demonstrated modest performance gains. Moreover, the amount of available training data was largely sufficient to capture bin-to-bin dissimilarities, with the kNN approach benefitting from a small degree of “regularization” afforded by the annotation-based dissimilarity in our multi-distance learning approach, as indicated by the small value of *α* selected for the final models and the modest difference in MAE relative to *α* = 0.

**Fig. 2.**
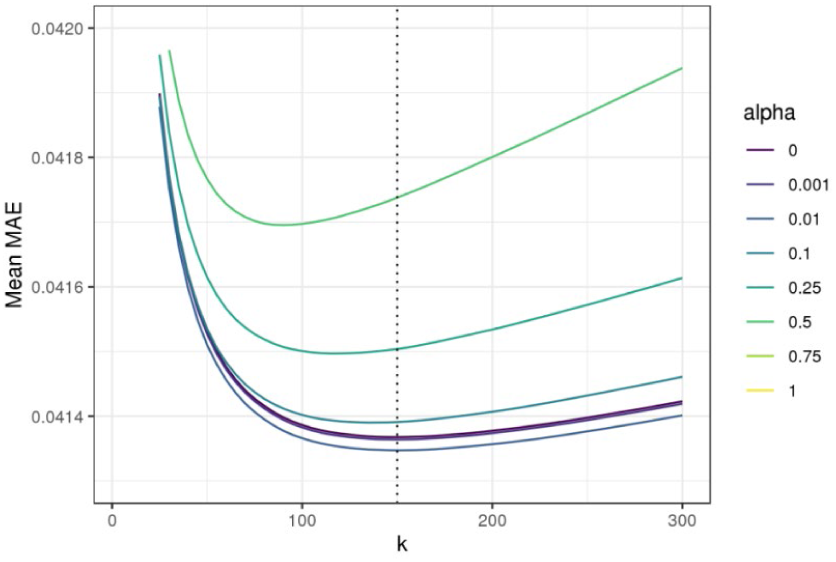
Cross-validated MAE measures for unweighted kNN approach. Figure is zoomed in to demonstrate separation across alpha values for best performing settings. Vertical dotted line indicates optimal *k* for *α* = 0.01.

To explore conditions of substantially reduced availability of training data, we additionally performed the same kNN model training with a random subset of 100 non-pregnant female samples. The optimal tuning parameter solution was *k* = 155 and *α* = 0.1 (MAE = 0.04298), in contrast to MAE = 0.04315 for *α* = 0.01 (selected from training on the complete data). The results of this simple sensitivity analysis follow intuition that a larger value of *α* would be selected in model training when more limited information available in ***X***.

### 3.2. Coverage correction

There were 476 negative trisomy NIPS samples available for comparative performance analysis. Using the wkNN with tuning parameters selected via the cross-validation results in Section 3.1, we derived corrected coverage profiles and compared them to results based on the annotation-based GAM model. Overall, we observed a median single-sample MAD reduction of 38.7% relative to the GAM regression approach, with the sample MAD consistently lower using our proposed wkNN method (range in MAD reduction: 26.5-53.5%). Further summary statistics of these results are presented in Table 1.

**Table 1.** Profile coverage correction MAD summaries for GAM and wkNN methods.

| MAD (N = 476) | GAM Model | wkNN Model | % Reduction |
| --- | --- | --- | --- |
| Minimum | 0.072 | 0.041 | 26.5 |
| 1<sup>st</sup> Quartile | 0.081 | 0.049 | 36.3 |
| Mean | 0.085 | 0.051 | 38.7 |
| Median | 0.085 | 0.052 | 38.7 |
| 3<sup>rd</sup> Quartile | 0.089 | 0.055 | 41.4 |
| Maximum | 0.118 | 0.075 | 53.5 |

Comparison of individual coverage profiles also demonstrated substantial smoothing effects of both variability and bias relative to the GAM approach. An illustrated example of typical coverage profile improvement (MAD reduction: 34.2%) is presented in Figure 3 for the trisomy prone chromosomes 13, 18 and 21.

**Fig. 3.**
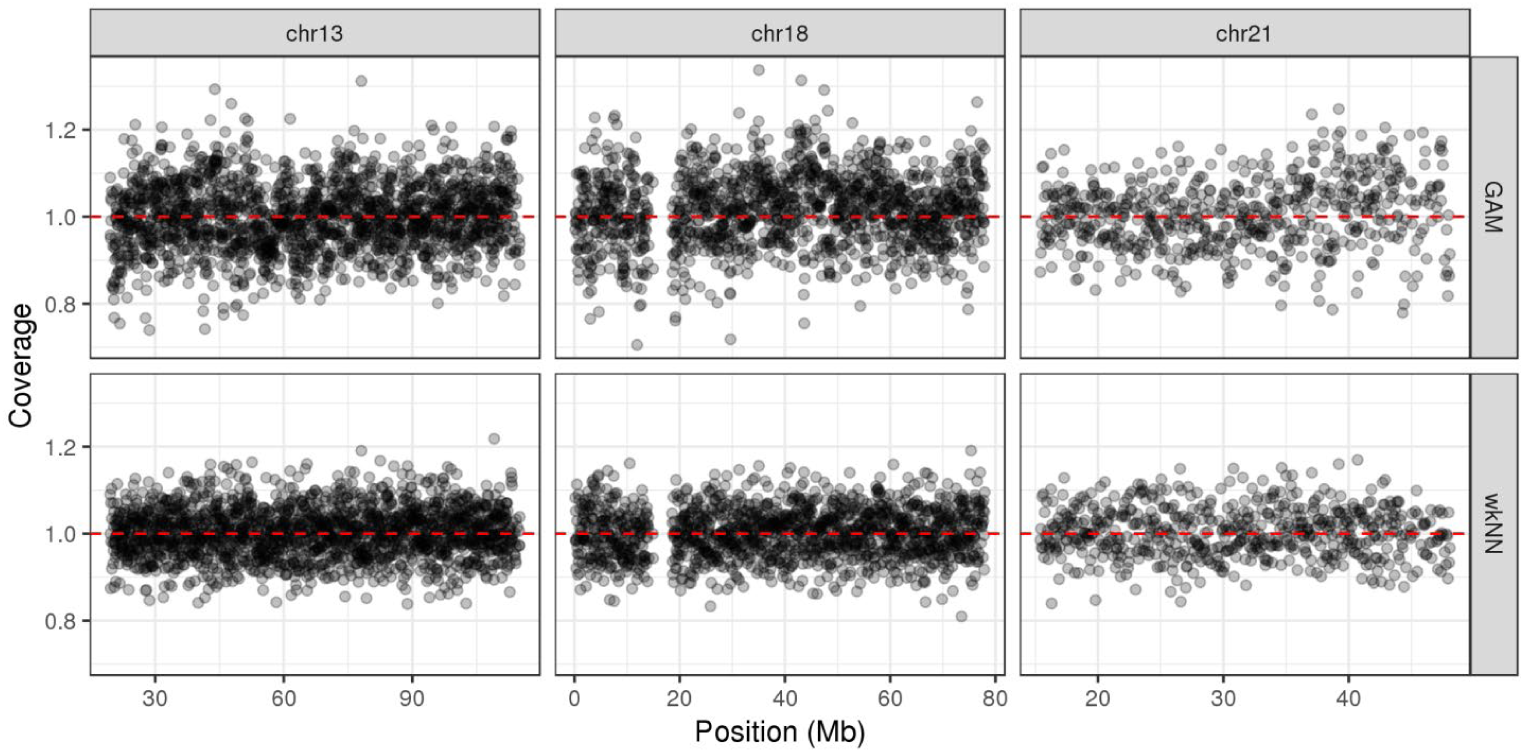
Corrected 50 kb bin coverage profiles using the GAM approach (top) vs. the proposed wkNN model (bottom) for an example negative trisomy NIPS test sample for chromosomes 13, 18, and 21 (columns).

### 3.3. Trisomy detection

A total of 300 negative trisomy NIPS samples were randomly selected to serve as a reference set for deriving estimates of *μ*_*c*_ and *SD*_*c*_ for *c* = 13,18,21. Corresponding Z-scores were generated for 176 negative trisomy samples and the 141 positive trisomy samples using chromsome proportions derived from corrected coverage profiles under each method (GAM vs wkNN). Boxplots of the Z-scores for the three chromosomes by trisomy status and the method of coverage profile used are depicted in Figure 4. Trisomy Z-scores among negative samples were highly correlated across method (Pearson *ρ*>0.69 for each chromosome), while signed-rank testing for increased positive case Z-scores were significant when assessed separately for all three chromosomes (all p-values < 0.0005). Results for trisomy 13 demonstrated the most substantial improvement, with a median Z-score increase of 7.08 among positive cases, and only four positive trisomy case Z-scores were higher for the GAM corrected coverage profiles relative to the wkNN approach. None of these four cases would have resulted in different NIPS results (all *Z*>6.0).

**Fig. 4.**
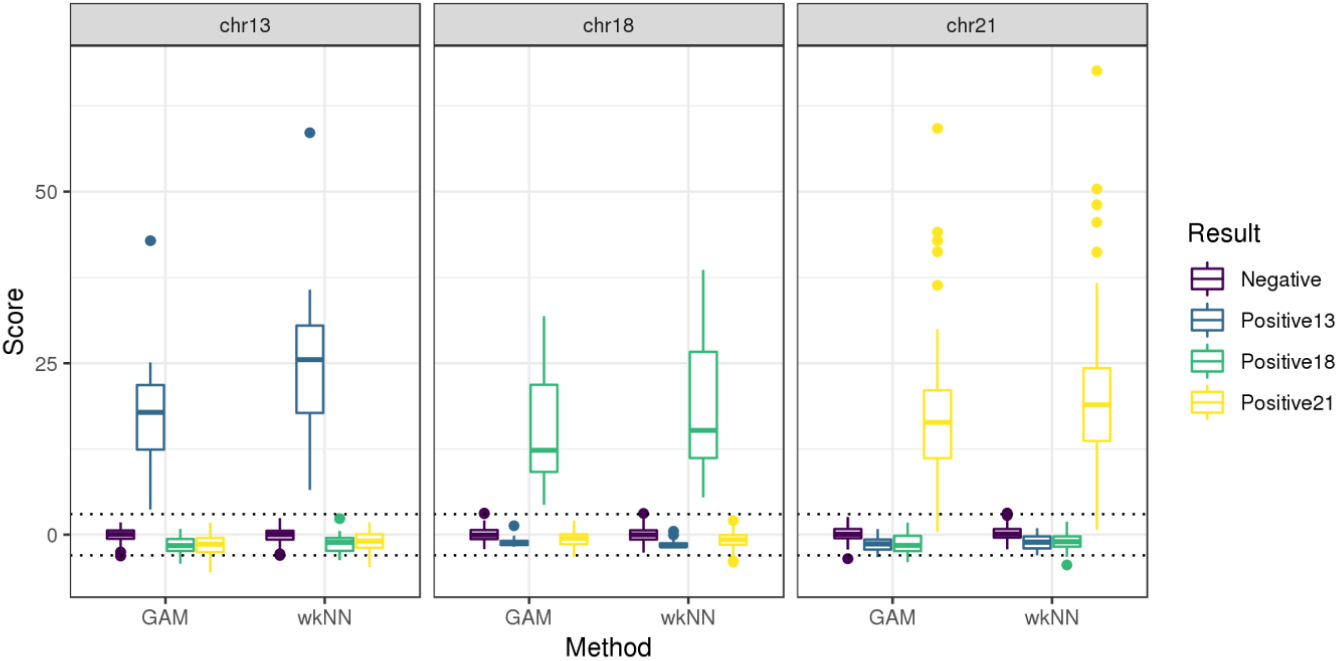
Boxplots of Z-scores used for trisomy screening, separated by method (GAM vs wkNN), chromosome, and trisomy status. Typical screening threshold values of (−3,3) are depicted by dotted horizontal lines.

## 4. Discussion

In this paper, we proposed a machine learning strategy to improve within-sample coverage profile correction for shallow-depth ccfDNA WGS. The noted inter- and intra-sample heterogeneity of these coverage profiles make the identification of even large structural alterations challenging in applications where the mutation frequency is anticipated to be low. Adopting similar principles to traditional methods based on GC-content correction through genomic bin stratification, we defined straight-forward and intuitive kNN-type approach to leverage empirical and annotation-based measures of genomic bin dissimilarity data under a multi-distance learning framework. In contrast to standard GC-content correction procedures implemented via GAM methods, our approach allows for the control of known and latent sources of coverage variability, using genomic annotation as a manner of regularizing empirical measures of dissimilarity in retrospective data.

Relative to the regression-based coverage profile correction method based solely on genomic annotation, our wKNN approach demonstrated substantial improvement in both coverage profile variability reduction and improved detection of positive fetal trisomies. Although overall screening performance was already nearly perfect, the increased discrimination distance between positive and negative cases indicates improved trisomy sensitivity and specificity at lower fetal fractions. Thus, adopting our coverage correction procedure should render the NIPS assay more robust to random fluctuations in sample fetal fractions. Our approach also makes the identification of micro-duplications and deletions more feasible. Similar methods could also be extended to capture-based assays for ccfDNA, given a sufficient number of capture regions.

A number of limitations and potential extensions warrant mention. The data used to derive our models were from healthy female donor plasma, which obviates the ability to provide coverage profile correction for chromosome Y. For NIPS, we could also universally exclude trisomy-prone chromosome from potential neighbors to make the approach more robust for trisomy detection. Alternative machine learning algorithms may also provide more accurate coverage correction performance if they were trained for each individual bin, although this would require fitting and validating *B* separate models. The grid search over values of *α* was fairly crude, and additional research improving how to tune *α* could lead to improved coverage profile correction. Similarly, we only considered two feature spaces for this particular multi-distance learning application, and extensions to ≥ 2 information sources (requiring *α* be defined as a simplex) would greatly complicate model training. Selected genomic bin size may heavily influence overall performance, particularly if smaller bins (e.g., 5-10 kb) are used instead of 50 kb. Under these conditions, our multi-distance learning approach may be particularly useful given the substantial increase in the bin dimensionality. Finally, it’s not clear how generalizable our approach is to external labs, where differences in sequencing conditions may yield different inter-bin correlation patterns. Further investigation into genomic annotations shared by neighboring bins may elucidate characteristics that contribute to the observed coverage biases of ccfDNA WGS.

## 5. Acknowledgments

Funding for this project was supported by the Mayo Clinic Center for Individualized Medicine.

